# Autophagy restricts fungal accommodation in the roots of *Arabidopsis thaliana*

**DOI:** 10.1101/2023.07.21.550010

**Authors:** Patricia Zecua-Ramirez, Ernesto Llamas, Nyasha M. Charura, Nick Dunken, Concetta De Quattro, Alexander Mandel, Gregor Langen, Yasin Dagdas, Alga Zuccaro

## Abstract

Endophytic colonization of *Arabidopsis thaliana* by the beneficial root endophyte *Serendipita indica* is characterized by an initial biotrophic phase followed by a restricted host cell death-associated phase. This latter phase involves regulated cell death (RCD) for fungal accommodation. However, the host molecular pathways that limit *S. indica* colonization and govern symbiosis remain largely unknown. Our study demonstrates that autophagy, a major cellular degradation pathway, is activated during *S. indica* colonization and is required to restrict fungal colonization in Arabidopsis. Independent Arabidopsis knockout (KO) mutants deficient in autophagosome formation are more susceptible to deoxyadenosine (dAdo), a cell death inducer produced by two secreted *S. indica* effectors at the onset of the cell death-associated phase. In the *atg5* autophagy mutant background, impaired dAdo uptake prevents dAdo-induced and symbiosis-mediated cell death. Based on our data, we propose that autophagy-mediated pro-survival responses in the host are crucial for maintaining a balanced symbiotic interaction between *S. indica* and Arabidopsis.

**In a Nutshell:** Our study reveals that during colonization of *Arabidopsis thaliana* roots by the beneficial root endophyte *Serendipita indica*, autophagy, a key cellular degradation pathway, is activated to limit fungal colonization. Autophagy-deficient Arabidopsis mutants are more susceptible to deoxyadenosine (dAdo), a cell death inducer produced by *S. indica*. We propose that autophagy-mediated pro-survival responses are essential for maintaining a balanced symbiotic interaction between *S. indica* and Arabidopsis.

## Introduction

Plant-colonizing fungi use diverse strategies to interact with hosts according to their lifestyles (Lo Presti et al., 2015). Beneficial root endophytic fungi of the order Sebacinales (Basidiomycota) establish mutualistic relationships with a variety of plant species (Weiss et al., 2016). These endophytic root associations result in growth promotion as well as enhanced resistance and tolerance against biotic and abiotic stresses (Oberwinkler et al., 2013; Weiss et al., 2016). To colonize plants, Sebacinales have evolved mechanisms to counteract the plant immune response and modulate the host metabolism, as well as processes involved in cell death (Deshmukh et al., 2006; Schafer et al., 2009; Jacobs et al., 2011; Nizam et al., 2019; Dunken et al., 2023).

Endophytic root colonization of *Arabidopsis thaliana* (hereafter Arabidopsis) and *Hordeum vulgare* (hereafter barley) by the beneficial fungus *Serendipita indica* (syn. *Piriformospora indica*) follows a biphasic strategy. *S. indica* establishes an initial biotrophic phase with hyphae enveloped by the host plasma membrane, followed by a cell death-associated phase restricted to the epidermis and cortex layers (Deshmukh et al., 2006; Qiang et al., 2012; Lahrmann et al., 2013). The latter is associated with the secretion of hydrolytic fungal enzymes (Zuccaro et al., 2011; Lahrmann and Zuccaro, 2012; Lahrmann et al., 2013; Lahrmann et al., 2015). The occurrence of this regulated cell death is thought to facilitate niche differentiation and ensures nutrient availability, without compromising plant growth (Deshmukh et al., 2006; Zuccaro et al., 2011; Weiss et al., 2016).

The combined activity of two effector enzymes secreted by *S. indica* during Arabidopsis and barley root colonization (Thurich et al., 2018; Nizam et al., 2019) was found to produce the DNA-derived active molecule deoxyadenosine (dAdo) (Dunken et al., 2023). Extracellular dAdo is imported into the cytoplasm by the Arabidopsis equilibrative nucleoside transporter 3 (ENT3), which functions as a transporter for purine and pyrimidine nucleosides such as adenosine and uridine (Li et al., 2003). dAdo has been previously described as a cell death inducer in animal systems (Thammavongsa et al., 2013; Winstel et al., 2018). In Arabidopsis, dAdo has been shown to trigger cell death, the accumulation of metabolites involved in stress signaling, and the upregulation of cell death marker genes (Dunken et al., 2023). This suggests that dAdo-mediated cell death contributes to the regulated and restricted symbiotic cell death observed with *S. indica*.

The induction of restricted cell death in host plants by *S. indica* is a crucial process for proper fungal accommodation. However, the host molecular mechanisms that limit colonization and allow the maintenance of a symbiotic relationship are largely unknown. Fungal colonization triggers a metabolic reprogramming of the host tissue, promoting the activation of pathways aimed at restoring homeostasis and restraining cell death. One potential host mechanism that could contribute to the regulation of this beneficial interaction is autophagy, a major degradation and nutrient recycling pathway in eukaryotes. Autophagy is activated by environmental and developmental stimuli and plays a vital role in maintaining cellular and metabolic homeostasis (He and Klionsky, 2009).

Autophagy is also known for its involvement in host immune responses and has been studied in various plant-microbe interactions (Hofius et al., 2017; Üstün et al., 2017; Leary et al., 2019). During biotrophic infections by pathogenic microbes, autophagy regulates the hypersensitive response (HR), which is a localized form of regulated cell death activated by intracellular immune receptors (Jones and Dangl, 2006; Coll et al., 2011). Autophagy components have been shown to promote HR upon infection with avirulent strains, depending on the nature of the immune receptor involved and the immune signaling pathway (Hofius et al., 2009; Hackenberg et al., 2013; Coll et al., 2014; Han et al., 2015; Munch et al., 2015). In non-infected tissue, autophagy plays a protective role by counteracting reactive oxygen species (ROS) production, salicylic acid (SA) signaling, endoplasmic reticulum (ER) stress, and promoting degradation of ubiquitinated proteins, derived from infection-induced systemic responses (Yoshimoto et al., 2009; Munch et al., 2014). During infections with necrotrophic fungi, autophagy promotes resistance and restricts disease-associated lesion formation to the site of infection (Lai et al., 2011; Lenz et al., 2011; Katsiarimpa et al., 2013). Additionally, components of the autophagy machinery participate in mutualistic interactions of *Phaseolus vulgaris* with rhizobacteria and arbuscular mycorrhizal fungi (Estrada-Navarrete et al., 2016).

In animals and plants, autophagy belongs to the protein homeostasis (proteostasis) network and mediates the lysosomal / vacuolar degradation of cytosolic damaged proteins, dysfunctional organelles and protein aggregates. In plants the autophagic cargo is engulfed in double-membrane vesicles called autophagosomes and delivered to the vacuole for breakdown and recycling (He and Klionsky, 2009; Marshall and Vierstra, 2018). Autophagy is broadly divided into microautophagy and macroautophagy (Liu and Bassham, 2012). Macroautophagy (hereafter referred to as autophagy) is mediated by proteins encoded by autophagy-related (ATG) genes. ATG proteins collectively coordinate the biogenesis of autophagosomes (Marshall and Vierstra, 2018).

To investigate whether autophagy is involved in *S. indica* colonization and host cell death, we employed Arabidopsis knockout (KO) mutants deficient in autophagosome formation, resulting in impaired autophagy. Here we show that these mutants exhibit increased fungal colonization during the cell death-associated phase, indicating that autophagy restricts colonization. Live-cell imaging revealed autophagosome formation in proximity to *S. indica* hyphae, indicating activation of autophagy at the microbe-host interface and suggesting a potential localized response to *S. indica* colonization. Additionally, autophagy mutants show increased sensitivity to dAdo, and their recovery after dAdo treatment is impaired, indicating that a functional autophagy pathway is essential to restore homeostasis and restrict cell death. Furthermore, in the autophagy mutant background, additional impairment of dAdo uptake confers resistance to dAdo-induced cell death and prevents the symbiosis-mediated cell death. Taken together, our results highlight the pro-survival role of autophagy in the beneficial interaction between *S. indica* and Arabidopsis. This shows that autophagy is a mechanism that restricts *S. indica* colonization and the extent of cell death, thereby maintaining a balanced symbiotic colonization.

## Results

### Autophagy is activated in response to *S. indica* and restricts fungal colonization

To determine whether host autophagy is involved in *S. indica* colonization, we quantified endophytic fungal colonization in the Arabidopsis autophagy KO mutants *atg5-3* and *atg10-1* using quantitative reverse transcription PCR (RT-qPCR). Before RNA extraction, the root colonized material was carefully washed to remove extraradical hyphae. ATG10 mediates the conjugation of ATG12 to ATG5, resulting in the formation of the ATG12-ATG5 complex. This complex, along with ATG16 promotes ATG8 lipidation, a key process for autophagic vesicle assembly. Mutation of ATG5 or ATG10 leads to impaired formation of autophagosomes (Thompson et al., 2005; Phillips et al., 2008). Both autophagy mutants exhibited significantly higher colonization levels than the Col-0 wild type (WT) at 6 days post-inoculation (dpi), which marks the onset of the cell death-associated phase (Fig. 1A). This suggests that fungal colonization is negatively regulated by host autophagy activation. Additionally, we observed an increase in the expression of the *S. indica*-induced marker gene AT1G58420, which is also responsive to ATP, dAdo, and wounding (Choi et al., 2014; Nizam et al., 2019; Dunken et al., 2023), in colonized roots of autophagy mutants compared to WT at 6 dpi (Fig. 1B). Colonization of *atg5-3* and *atg10-1* roots also led to an enhanced expression of the immune-related marker genes *WRKY33* and *CYP81F2* compared to WT (Fig. S1), which are involved in the biosynthesis of secondary antimicrobial compounds (Bednarek et al., 2009; Clay et al., 2009; Birkenbihl et al., 2017).

**Figure 1:**
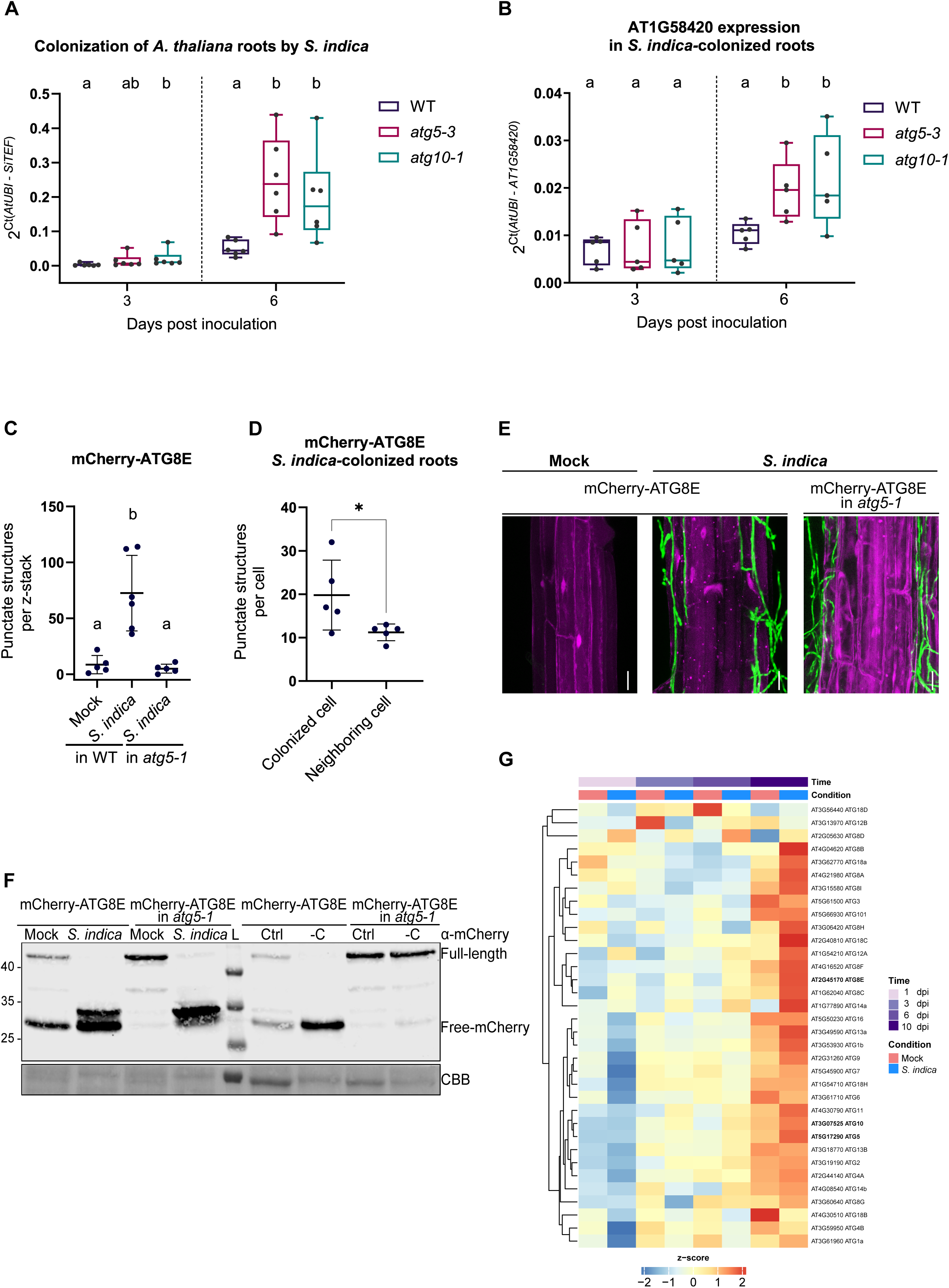
Involvement of autophagy in the colonization of *A. thaliana* roots by *S. indica*. (A) *S. indica* colonization of WT and autophagy mutants *atg5-3* and *atg10-1* quantified by RT-qPCR at 3 and 6 days post inoculation (dpi). Fungal colonization in plant root tissue was calculated from the ratio of *S. indica* (*SiTEF*) to plant (*AtUbi*) using cDNA as template and the 2^−ΔCt^ method. Boxplots with whiskers extending to the minimum and maximum values represent data from 6 independent biological replicates. Different letters indicate significant differences (p < 0.05) according to Kruskal-Wallis test and post-hoc Dunn test using Benjamini-Hochberg for false discovery rate correction. (B) Expression of the marker gene AT1G58420 in *S. indica*-colonized WT, *atg5-3* and *atg10-1* roots at 6 dpi. Relative expression was calculated compared to plant ( *AtUbi*) using cDNA as template and the 2^−ΔCt^ method. Boxplots with whiskers extending to the minimum and maximum values represent data from 5 independent biological replicates. Different letters indicate significant differences (p < 0.05) according to Kruskal-Wallis test and post-hoc Dunn test using Benjamini-Hochberg for false discovery rate correction. (**C**) Quantification of mCherry-ATG8E-labeled puncta was performed using confocal images of transgenic roots expressing mCherry-ATG8E. Confocal microscopy was conducted on differentiated root cells following mock or *S. indica* inoculation. The plot (mean *±* SD) shows the total number of puncta per z-stack in the whole 63X captured image for mCherry-ATG8E mock (n=5), *S. indica* (n=6) in WT and mCherry-ATG8E in *atg5-1* background *S. indica* (n=5). Different letters indicate significant differences (p < 0.05) according to Kruskal-Wallis test and post-hoc Dunn test using Benjamini-Hochberg for false discovery rate correction. (**D**) Quantification of mCherry-ATG8E-labeled puncta on differentiated root cells colonized by *S. indica*. The plot (mean *±* SD) shows the number of mCherry-ATG8 puncta per root cell distinguishing between cells with hyphal contact and neighboring cells without hyphal contact (n=5). The asterisk indicates significant differences (p < 0.05) according to unpaired Student’s t-test. (**E**) Confocal laser scanning microscopy (CLSM) representative images obtained of transgenic roots expressing mCherry-ATG8E in WT or *atg5-1* background. The microscopy was conducted from the epidermis of differentiated root cells, following mock or *S. indica* treatment for 10 days after seed inoculation. The images represent maximal projections of 8-10 optical sections, scale bar 20 µm. (**F**) Autophagy flux analysis was performed using transgenic plants expressing mCherry-ATG8E in WT or *atg5-1* background. The left panel displays a blot with samples obtained from collected roots upon mock or *S. indica* treatment for 10 days after seed inoculation. The right panel displays a blot with samples obtained from collected seedlings incubated in control or carbon-depleted media for starvation-induced autophagy. Immunoblot analysis was performed using an anti-mCherry antibody. Coomassie Brilliant Blue (CBB) was used as protein loading control. (**G**) Heatmap with expression levels of autophagy-related genes (ATG) from Arabidopsis WT roots samples collected at 1, 3, 6 and 10 days post inoculation with mock or *S. indica*. Genes with an average TPM value > 1 TPM across all samples were selected. The heatmap shows the z-score of log2 transformed TPM + 1 values of selected Arabidopsis genes. For each condition the average expression across the three biological replicates is represented. The full version of the heatmap of autophagy-associated genes can be found in Figure S2.

To monitor autophagy activation during *S. indica* colonization, we used Arabidopsis transgenic lines expressing *pUbi::mCherry-ATG8E* in the WT background or in the *atg5-1* KO mutant background, serving as a control. ATG8 is commonly used as a marker to detect and quantify autophagy levels (Yoshimoto et al., 2004; Liu and Bassham, 2012; Klionsky et al., 2021). Confocal microscopy in the differentiation zone of mCherry-ATG8E WT colonized roots revealed an increased amount of mCherry-ATG8E-labelled punctate structures, resembling autophagosomes, compared to mock-inoculated plants (Fig. 1C, E). In the *atg5-1* mutants, colonized roots exhibited fewer structures with similarity to the larger aggregates previously observed in autophagy-defective mutants (Li et al., 2015) (Fig. 1C, E). We then focused our attention on colonized roots of mCherry-ATG8E in the WT background, where we observed that cells in contact with *S. indica* hyphae exhibited an increased number of punctate structures compared to the nearby cells lacking hyphal contact (Fig. 1D, E). These findings provide evidence that *S. indica* induces localized autophagy at the site of interaction.

We also examined autophagy activation in mock and *S. indica*-colonized mCherry-ATG8E root tissue by immunoblotting. The autophagic turnover of mCherry-ATG8E labelled autophagosomes results in accumulation of the free fluorescent mCherry protein in the vacuole due to faster decay of the ATG8 segment (Chung et al., 2010). Colonization by *S. indica* led to an increased proportion of free mCherry compared to mock treatment, confirming induction of autophagic activity by the fungus (Fig. 1F). Interestingly, we detected an additional mCherry-ATG8E cleavage fragment produced in both WT and mutant background in the presence of *S. indica*, independent from the *atg5-1* mutation. This additional fragment is not produced upon autophagy induction due to carbon starvation (Fig. 1F). These findings suggest that, in addition to the induction of autophagy, *S. indica* colonization triggers an ATG5-independent hydrolysis of ATG8. Furthermore, we analyzed transcriptomic data from Arabidopsis roots colonized by *S. indica* at different time points. We observed an induction in the expression of autophagy-associated genes including the autophagy-related genes (ATGs) of the core autophagy machinery (Mizushima et al., 2011; Liu and Bassham, 2012) (Fig. 1G, S2). Taken together, these results demonstrate that autophagy is locally activated during *S. indica* colonization regulating fungal accommodation in the roots of Arabidopsis.

### Autophagy promotes cell survival during dAdo-induced cell death

During the cell death-associated phase of *S. indica* colonization, two fungal apoplastic effector enzymes produce extracellular dAdo. dAdo induces cell death in various plant species, including Arabidopsis (Dunken et al., 2023). Since autophagy plays a role in promoting cell survival and maintenance of cellular homeostasis, we investigated whether it could restrict dAdo-induced cell death. To test this hypothesis, we measured the onset of cell death by monitoring the photosynthetic efficiency, which is a key indicator of plant health (Dunken et al., 2022). Our results showed that treatment with dAdo strongly decreased the maximum quantum yield of photosystem II (F_V_/F_M_) in *atg5-3* and *atg10-1* KO mutants compared to WT, leading to earlier cell death (Fig. 2A, C and S3). Similar enhanced sensitivity upon treatment with dAdo was observed during seed germination tests with *atg5-3* mutants (Fig. S4). As a positive control, we additionally assessed the sensitivity of these mutants to methyl jasmonate (MeJA), a known inducer of senescence (He et al., 2002). The results showed that autophagy mutants exhibited increased sensitivity to MeJA (Fig. 2B, C and S3). We further investigated the role of autophagy in dAdo-induced cell death by testing other mutants of the autophagic machinery. Treatment with dAdo affected the germination of *atg2-2* seeds and significantly reduced the photosynthetic efficiency of *atg11-1* compared to WT (Fig. S4, S5). The effects of dAdo on photosynthetic activity have been shown to be concentration-dependent (Dunken et al., 2023). Similarly, autophagy mutants exhibited a dose-dependent response, indicating a regulated sensitivity to dAdo (Fig. S6). These findings further support the pro-survival role of autophagy in response to stress and cell death.

**Figure 2:**
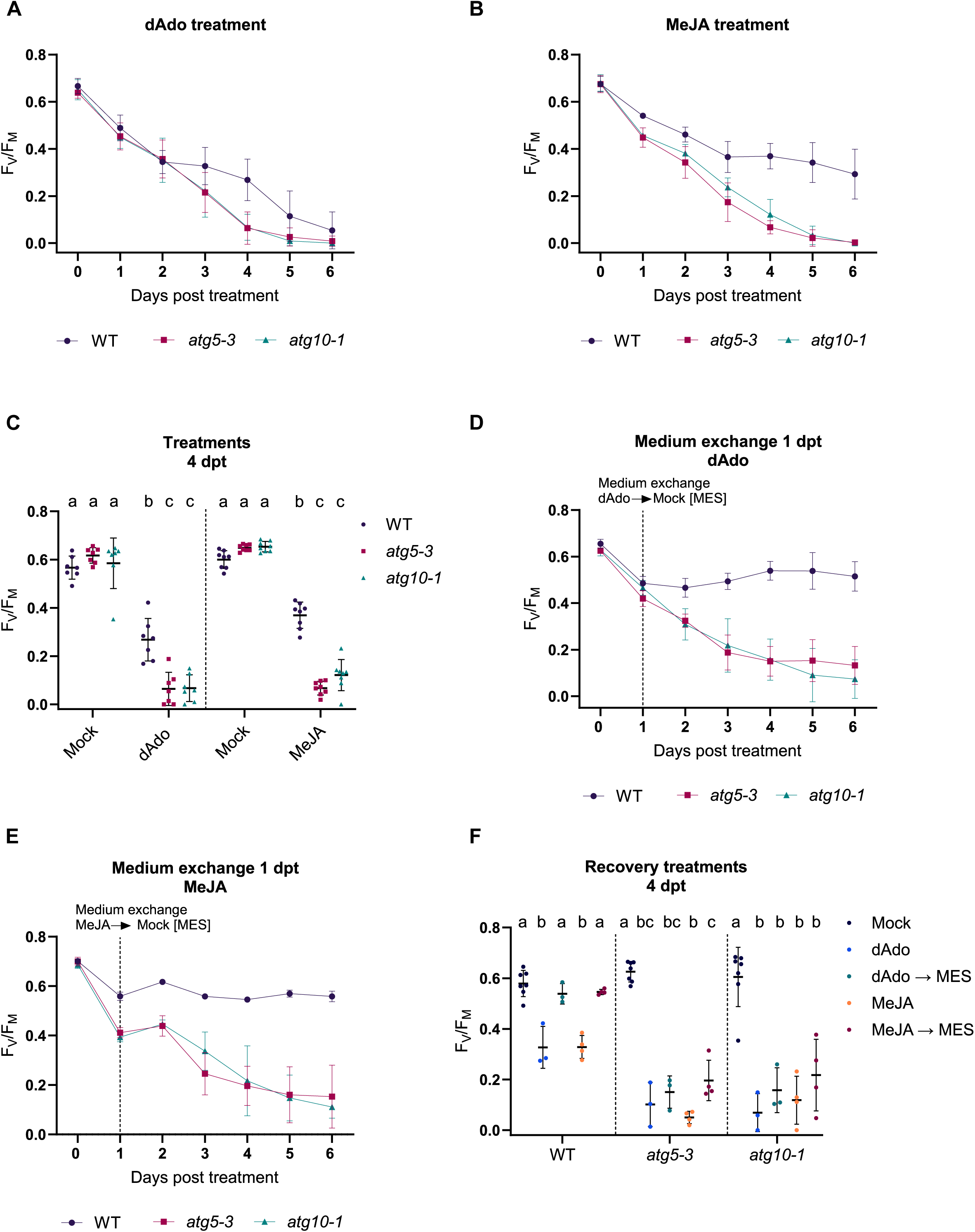
Autophagy promotes cell survival during dAdo-induced cell death. (**A**) Photosystem II maximum quantum yield (F_V_/F_M_) of nine-day-old seedlings treated with mock (MES 2.5 mM buffer) or dAdo (500 µM), measured by PAM fluorometry. Error bars represent *±* SD of the mean of 7 independent biological replicates. (**B**) Photosystem II maximum quantum yield (F_V_/F_M_) of nine-day-old seedlings treated with mock (MES 2.5 mM buffer) or methyl jasmonate (MeJA, 500 µM) as positive control for cell death, measured by PAM fluorometry. Error bars represent *±* SD of the mean of 8 independent biological replicates. (**C**) Quantification of F_V_/F_M_ of WT and autophagy mutants *atg5-3* and *atg10-1*, 4 days post treatment (dpt). The plot (mean *±* SD) represents data from 7-8 independent biological replicates, each consisting of 12 wells with 3 seedlings per well. Different letters indicate significant differences (p < 0.05) according to one-way ANOVA with post-hoc Tukey HSD test. (**D**) Photosystem II maximum quantum yield (F_V_/F_M_) of nine-day-old seedlings treated with mock (MES 2.5 mM buffer) or dAdo (500 µM), measured by PAM fluorometry. dAdo treatment solution was replaced after 24 h with MES 2.5 mM buffer. Error bars represent *±* SD of the mean of 3 independent biological replicates. (**E**) Photosystem II maximum quantum yield (F_V_/F_M_) of nine-day-old seedlings treated with mock (MES 2.5 mM buffer) or methyl jasmonate (MeJA, 500 µM), measured by PAM fluorometry. MeJA treatment solution was replaced after 24 h with MES 2.5 mM buffer. Error bars represent *±* SD of the mean of 4 independent biological replicates. (**F**) Quantification of F_V_/F_M_ of WT and autophagy mutants *atg5-3* and *atg10-1*, 4 days after treatment solution replacement. The plot (mean *±* SD) represents data from 3-4 independent biological replicates, each consisting of 12 wells with 3 seedlings per well. Different letters indicate significant differences (p < 0.05) according to one-way ANOVA with post-hoc Tukey HSD test.

To study the protective role of autophagy in more detail, we tested the ability of autophagy mutants to recover after dAdo and MeJA treatment by monitoring photosynthetic efficiency. We replaced dAdo and MeJA treatment solutions with buffer at 24 hours post-treatment. While WT seedlings recovered, the autophagy *atg5-3* and *atg10-1* mutants did not recover from cell death induction after treatment with both dAdo (Fig. 2D, F) and MeJA (Fig. 2E, F). These observations suggest that impaired autophagy prevents the resolution of metabolic stress induced by the treatments, resulting in cell death. Notably, we observed that recovery was also impaired in WT seedlings after 48 h of treatment (Fig. S7), suggesting the occurrence of a point-of-no-return during the execution of dAdo-induced cell death. Collectively, these results support the notion that autophagy exerts a detoxifying function against cellular stress induced by dAdo.

### Impaired dAdo uptake in *atg5-3* mutant plants attenuates dAdo-induced and symbiosis-mediated cell death

The import of extracellular dAdo by the ENT3 transporter is required for dAdo-dependent regulated cell death (Dunken et al., 2023). We hypothesized that impaired dAdo uptake in *atg5-3* mutant plants would generate dAdo resistance, thereby reducing cell death. To test this effect, we generated an *atg5 ent3* double mutant. Examination of the photosynthetic efficiency in seedlings incubated with mock or dAdo showed that the double mutant gained resistance to dAdo, similar to the *ent3* mutant, exhibiting reduced sensitivity (Fig. 3A). The double mutant showed increased sensitivity to MeJA, indicating that mutation of the ENT3 transporter confers a specific resistance phenotype to the cell death trigger dAdo but not to MeJA (Fig. 3B). To evaluate the extent of cell death, we used Evans blue dye, which selectively permeates cells with ruptured plasma membranes (Vijayaraghavareddy et al., 2017). After a 4-day treatment with dAdo or mock solution, we quantified the stained dead cells at the root tip, where cell death was most pronounced, by using bright-field microscopy (Fig. 3C, D). The *atg5-3* autophagy mutant displayed enhanced cell death, while impairment of the ENT3 transporter abolished the cell death phenotype compared to WT. The *atg5 ent3* double mutant exhibited an intermediate dAdo-sensitive phenotype between *atg5-3* and *ent3* (Fig. 3C, D). These findings indicate that impaired dAdo uptake in *atg5 ent3* mutant plants leads to reduced cell death and increased resistance to dAdo.

**Figure 3:**
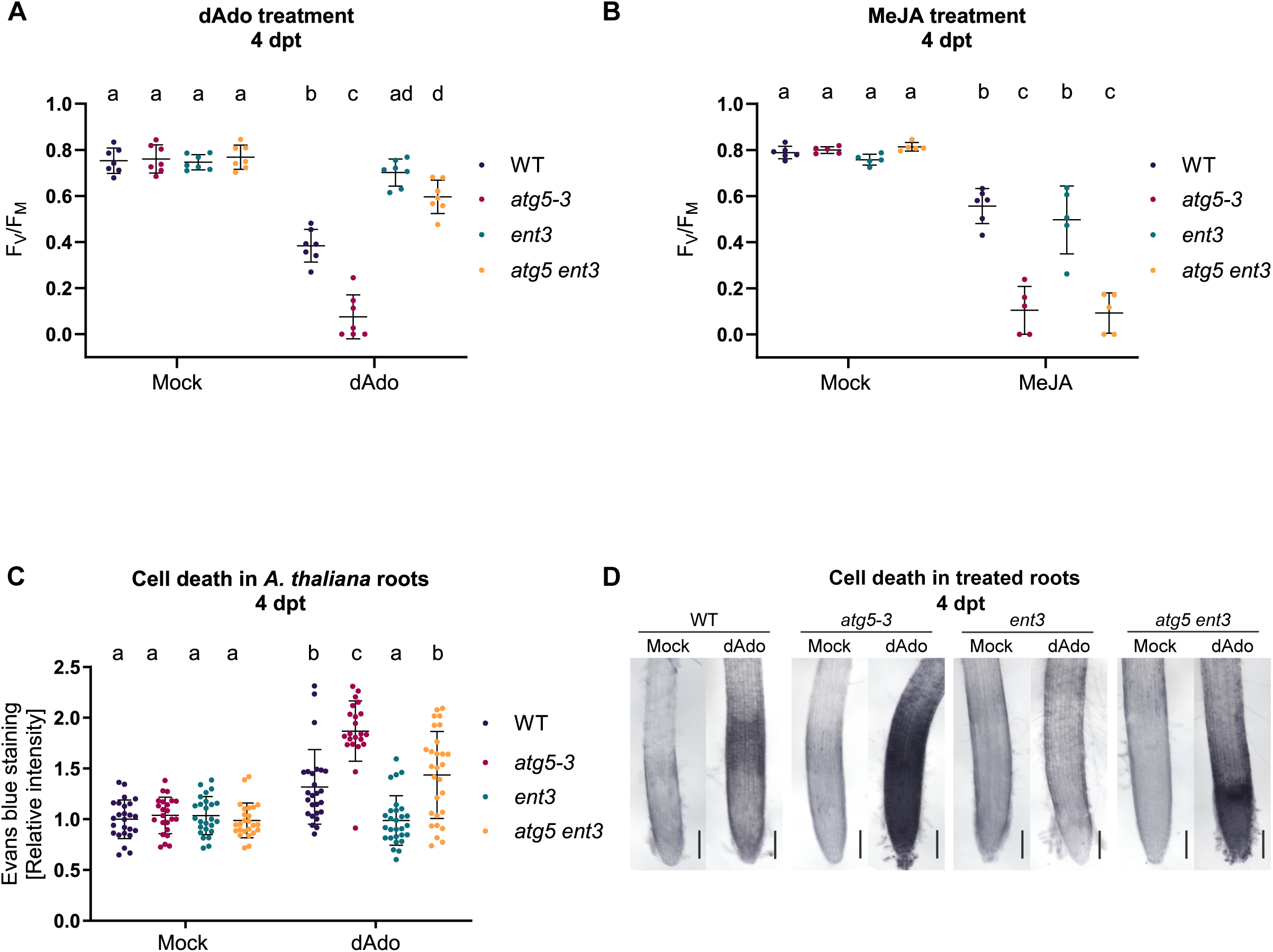
The double mutant *atg5 ent3* displays a dAdo resistance phenotype. (**A**) Quantification of photosystem II maximum quantum yield (F_V_/F_M_) of WT, *atg5-3*, *ent3* and *atg5 ent3* at 4 days post treatment (dpt) by PAM fluorometry. Nine-day-old seedlings were treated with mock (MES 2.5 mM buffer) or dAdo (500 µM). The plot (mean ± SD) represents data from 7 independent biological replicates, each consisting of 12 wells with 3 seedlings per well. Different letters indicate significant differences (p < 0.05) according to one-way ANOVA with post-hoc Tukey HSD test. (**B**) Quantification of photosystem II maximum quantum yield (F_V_/F_M_) of WT, *atg5-3*, *ent3* and *atg5 ent3* at 4 dpt by PAM fluorometry. Nine-day-old seedlings were treated with mock (MES 2.5 mM buffer) or methyl jasmonate (MeJA, 500 µM). The plot (mean ± SD) represents data from 5 independent biological replicates, each consisting of 12 wells with 3 seedlings per well. Different letters indicate significant differences (p < 0.05) according to one-way ANOVA with post-hoc Tukey HSD test. (**C**) Quantification of cell death at the root tip of WT, *atg5-3*, *ent3* and *atg5 ent3*. Eight-day-old seedlings were treated with mock (sterile milli-Q water) or dAdo (500 µM) and stained with Evans blue at 4 dpt. The plot (mean ± SD) represents relative values to WT mock from 22-28 biological replicates. Different letters indicate significant differences (p< 0.05) according to Kruskal-Wallis test and post-hoc Dunn test using Benjamini-Hochberg for false discovery rate correction. (**D**) Bright-field microscopy of root cell death at the root tip in WT, *atg5-3*, *ent3* and *atg5 ent3* seedlings incubated with mock or dAdo (500 µM) and stained with Evans blue dye at 4 dpt. Scale bar: 500 µm.

Symbiosis-mediated cell death by *S. indica* is supported by the production of dAdo (Dunken et al., 2023). To assess the impact of reduced sensitivity to dAdo in the *atg5 ent3* double mutant on root colonization, we performed colonization assays on ½ MS media and evaluated cell death using Evans blue dye at 12 dpi. Colonization by *S. indica* led to cell death in the differentiation zone of WT and *atg5-3* seedlings compared to mock conditions. Notably, *S. indica*-induced cell death was abolished in the *ent3* mutant, and the *atg5 ent3* double mutant did not exhibit a significant increase in cell death under *S. indica* treatment compared to mock conditions (Fig. 4A, B). This observation suggests that the cell death induced by *S. indica* is prevented in the *ent3* and *atg5 ent3* double mutant plants.

**Figure 4:**
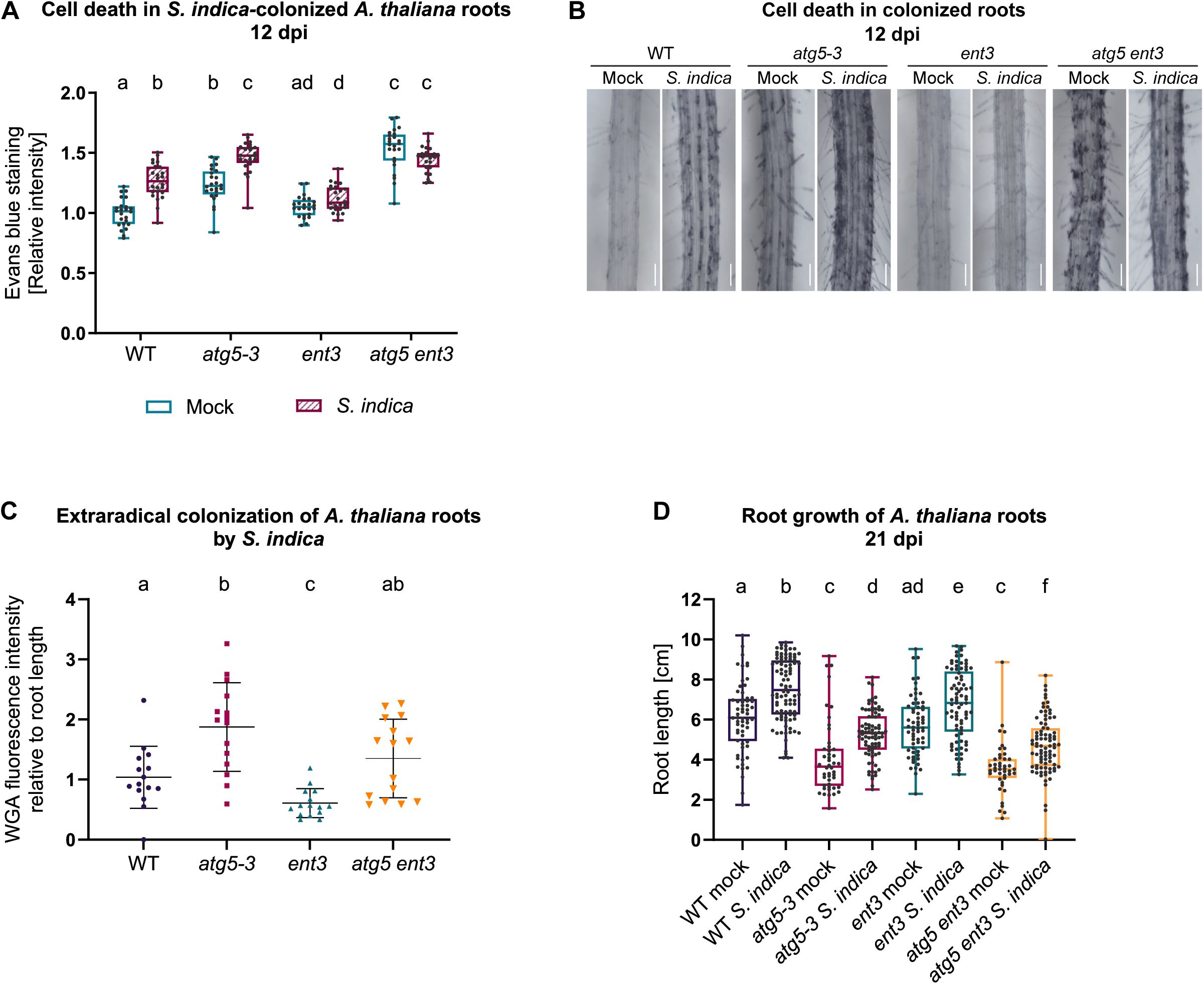
*S. indica*-mediated cell death is prevented in the double mutant *atg5 ent3*. (**A**) Quantification of cell death at the root differentiation zone in WT, *atg5-3*, *ent3* and *atg5 ent3*. Nine-day-old seedlings were inoculated with mock or *S. indica* and stained with Evans Blue dye at 12 days post inoculation (dpi). Boxplots with whiskers extending to the minimum and maximum values represent relative values to WT mock from 6 biological replicates. Different letters indicate significant differences (p < 0.05) according to Kruskal-Wallis test and post-hoc Dunn test using Benjamini-Hochberg for false discovery rate correction. (**B**) Bright-field microscopy of root cell death at the differentiation zone in WT, *atg5-3*, *ent3* and *atg5 ent3* seedlings inoculated with mock or *S. indica* and stained with Evans blue dye at 12 dpi. Scale bar: 500 µm. (**C**) Quantification of extraradical *S. indica* colonization on 1/10 PNM was assessed by fluorescence intensity using WGA-AF 488 staining 12 days after seed inoculation. The plot (mean ± SD) represents WGA fluorescent intensity values relative to root length and normalized to Col-0 mock, based on 15 biological replicates. Different letters indicate significant differences (p < 0.05) according to Kruskal-Wallis test and post-hoc Dunn test using Benjamini-Hochberg for false discovery rate correction. (**D**) Quantification of the primary root length of WT, *atg5-3*, *ent3* and *atg5 ent3* seedlings under mock or *S. indica* treatment, 21 days after seed inoculation. Boxplots with whiskers extending to the minimum and maximum values represent data from 43-92 biological replicates. Different letters indicate significant differences (p < 0.05) according to Kruskal-Wallis test and post-hoc Dunn test using Benjamini-Hochberg for false discovery rate correction.

To examine the impact of nutrient availability on autophagy, *S. indica* colonization and host growth, colonization experiments were performed using 1/10 PNM, a plant low-nutrient medium known to enhance the beneficial interaction between root endophytes and host plants (Lahrmann et al., 2013). Extraradical colonization was assessed using the chitin-binding lectin stain WGA-AF488, which revealed increased colonization in *atg5-3* compared to WT after 12 days (Fig. 4C, S8). At 21 days, we also observed increased endophytic colonization in both *atg5-3* and the *atg5 ent3* double mutant, as determined by RT-qPCR (Fig. S9). Mutation of ENT3 results in a transient reduction of the colonization at 12 dpi, which was not visible at a later stage (Fig. 4C, S8, S9; Dunken et al., 2023). This suggests that ENT3, and thus intracellular dAdo signaling, contributes to the onset of cell death but not after the endophyte is fully established in the roots of Arabidopsis. In contrast, autophagy remains important or becomes even more critical at later stages, likely playing a major role in maintaining a balanced colonization level.

To explore the involvement of autophagy in *S. indica*-mediated root growth, the primary root length of colonized seedlings was quantified after 21 days. Treatment with *S. indica* resulted in longer primary roots in WT plants compared to mock conditions. This increase in root growth was also observed in the *atg5-3* and *ent3 atg5* double mutant plants under these growth conditions (Fig. 4D), suggesting that impairment of autophagy does not interfere with the root growth phenotype induced by *S. indica*. Overall, our results endorse the activation of autophagy as a mechanism that restricts fungal accommodation, safeguards against dAdo-associated stress and cell death, and consequently contributes to maintaining a balanced interaction with *S. indica* (Fig. 5).

**Figure 5:**
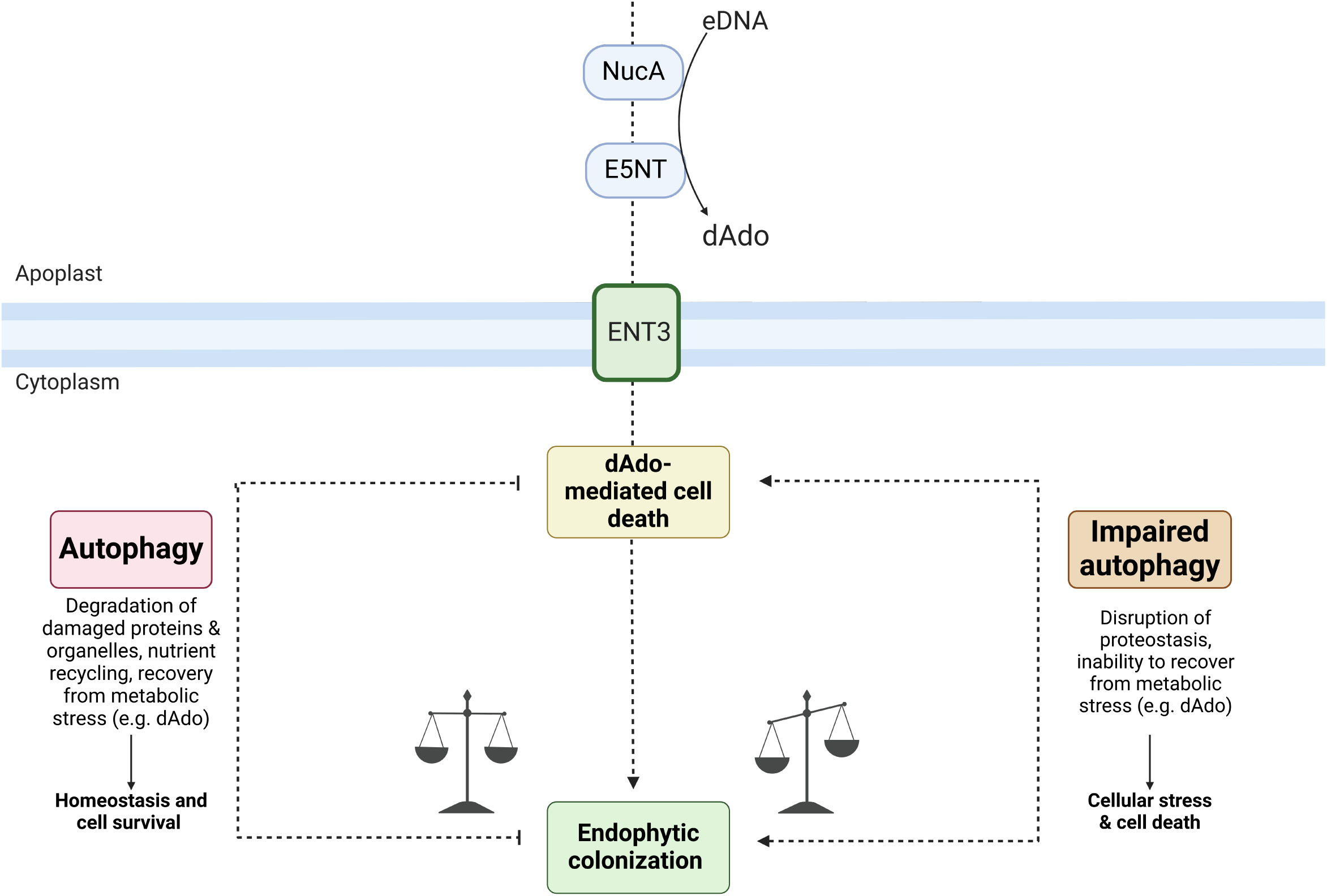
Current model for the crosstalk between autophagy and dAdo cell death during *S. indica* colonization. Endophytic root colonization of the host plant *Arabidopsis thaliana* by the beneficial root endophyte *Serendipita indica* (*S. indica*) is characterized by a biotrophic phase followed by a restricted cell-death associated phase. The activity of two secreted effector enzymes of *S. indica*, E5NT and NucA, produces a DNA-derived deoxynucleoside called deoxyadenosine (dAdo) (Dunken et al., 2023). Extracellular dAdo is imported into the cytoplasm by the Arabidopsis equilibrative nucleoside transporter ENT3, activating a dAdo-dependent regulated cell death. Autophagy, a major degradation and nutrient recycling pathway that maintains cellular homeostasis and modulates metabolism, plays a key role in the beneficial interaction between *S. indica* and Arabidopsis. Autophagy restricts fungal colonization and provides a protective function against dAdo-induced cell death. We propose that autophagy-driven pro-survival responses, such as the degradation of damaged proteins or protein aggregates, recycling of nutrients and recovery from metabolic stress facilitate a balanced symbiotic interaction. Illustration was designed using the Biorender online tool.

## Discussion

Our investigation of the interaction between the plant host Arabidopsis and the beneficial endophytic fungus *S. indica* highlights the role of autophagy in regulating fungal accommodation in the roots of this host. Our results show that autophagy is activated during colonization, particularly at later stages, suggesting a regulatory transcriptional mechanism triggered by fungal colonization. Loss of function in ATG5 and ATG10 leads to increased fungal colonization, especially during the cell death-associated phase and over the long term. Additionally, we analyzed the transcriptional response to *S. indica* in autophagy mutant roots. The higher expression of the *S. indica*-responsive marker gene AT1G58420, along with the transcription factor WRKY33 and its target CYP81F2 in colonized autophagy mutants, suggests a potential link between autophagy and the plant immune response. WRKY33 and CYP81F2 are involved in the production of secondary metabolites, such as glucosinolates, and play a role in regulating plant hormone pathways (Birkenbihl et al., 2017). This finding aligns with previous research demonstrating the interaction between WRKY33 and ATG18a during necrotrophic infections in Arabidopsis (Lai et al., 2011). However, the higher fungal biomass in the autophagy mutants could account for the increased expression of these immune-related marker genes.

To confirm autophagy activation in *S. indica*-colonized root cells, we examined autophagosome formation using mCherry-ATG8E transgenic lines. Our results showed the presence of structures resembling autophagosomes in colonized cells and to a lesser extent in neighboring cells, but not in the *atg5-1* line (Fig. 1C-E). The occurrence of autophagosomes during autophagy at the host-microbe interface has previously been observed as part of the localized immune response during pathogen infection (Dagdas et al., 2018; Pandey et al., 2021).

Immunoblot analysis showed increased mCherry-ATG8E cleavage, indicating higher autophagic flux during fungal colonization. The observed mCherry-ATG8E cleavage pattern in the *S. indica*-colonized *atg5-1* line (Fig. 1F) suggests the involvement of an independent mechanism of hydrolysis in addition to ATG5-dependent autophagy. This effect can be attributed to the hydrolytic activity occurring during the cell death-associated phase, where plant lytic enzymes, including proteases, may contribute to the degradation process. Additionally, this colonization stage is characterized by the release of numerous fungal proteases, which may also contribute to the observed band (Zuccaro et al., 2011; Lahrmann and Zuccaro, 2012; Lahrmann et al., 2015). However, it cannot be ruled out that other host degradation mechanisms may be activated in the presence of the fungus when the autophagic pathway is non-functional. One potential mechanism is the ubiquitin-proteasome system, which has been reported to have crosstalk with autophagy (Raffeiner et al., 2023).

During the symbiotic interaction, the fungus induces alterations in the host metabolic and transcriptional profiles. The uptake of dAdo initiates a cascade of events, characterized by the upregulation of cell death marker genes, electrolyte leakage, activation of the 26S proteasome, and extracellular accumulation of the retrograde stress signaling metabolite methylerythritol cyclodiphosphate (MEcPP) (Dunken et al., 2023). Our investigation reveals the involvement of autophagy in restricting dAdo-induced cell death. The increased sensitivity of *atg5-3*, *atg10-1*, *atg2-2*, and *atg11-1* mutants to dAdo, coupled with impaired recovery in *atg5-3* and *atg10-1* after dAdo removal (Fig. 2C, D), highlights the pro-survival function of autophagy. Notably, the additional mutation of ENT3 in the *atg5* background reduces cell death, conferring resistance to dAdo. This collective evidence supports the notion that dAdo-induced cell death in autophagy mutants is governed by a regulated mechanism.

Autophagy as a degradation pathway promotes a protective response during certain stress and cell death-related processes. For instance, autophagy prevents the precocious onset of senescence and facilitates nutrient remobilization during senescence (Schippers et al., 2007). Treatment with MeJA led to accelerated senescence-associated cell death in autophagy mutants (Fig. 2B, C), resembling the early senescence phenotype previously described for different autophagy mutants including *atg5* and *atg10* (Doelling et al., 2002; Hanaoka et al., 2002; Thompson et al., 2005; Phillips et al., 2008; Yoshimoto et al., 2009). Autophagic responses, known to manifest during senescence and various stress conditions, may potentially mitigate the impact of dAdo in a similar fashion. These protective functions involve the degradation of protein aggregates, recycling of damaged organelles (Xiong et al., 2007; Wada et al., 2009; Munch et al., 2014) and breakdown of pro-death signals (Hayward et al., 2009), contributing to the maintenance of proteostasis. Hence, we propose that the activation of autophagy could function as a regulatory mechanism providing support against dAdo-induced stress and limiting the extent of cell death.

Fungal-induced cell death mediated by apoplastic dAdo production and subsequent uptake via ENT3 contributes to the regulated *S. indica*-symbiotic cell death. Mutation of ENT3 impairs fungal-mediated cell death (Fig. 4A and Dunken et al., 2023). Consequently, the colonized *atg5 ent3* double mutant does not exhibit increased cell death mediated by the fungus (Fig. 4A). However, the double mutant displays a phenotype reminiscent of the *atg5-3* mutant, characterized by increased colonization levels (Fig. S9). Both *atg5* and the *atg5 ent3* double mutants exhibit higher levels of cell death in their roots, independent of *S. indica* presence, potentially creating favorable conditions for fungal colonization. This effect is more pronounced in the *atg5 ent3* double mutant due to the loss of function of the nucleoside transporter ENT3 (Fig. 4A). While the *ent3* mutant line shows no discernible phenotypic changes under mock conditions, the absence of autophagy in the double mutant leads to increased cellular stress. This suggests an interplay between autophagy and nucleotide metabolism. In plants and mammals, depletion of cellular purine nucleotide levels has been shown to trigger the activation of autophagy by inhibiting TOR (Target of Rapamycin), a key regulator of growth and nutrient sensing that negatively regulates autophagy (Hoxhaj et al., 2017; Kazibwe et al., 2020). Additionally, *At*ENT3 has been proposed to participate in the salvage pathway of nucleotide synthesis (Li et al., 2003). Therefore, alterations in nucleotide levels may increase cellular stress in the double mutant.

We also examined Arabidopsis root growth and confirmed that under nutrient-limiting conditions, *S. indica* induced primary root growth (Fig. 4D). This phenotype was also observed in the *atg5-3* mutant, suggesting that while autophagy influences fungal-mediated cell death and colonization, it does not affect the root growth phenotype. Collectively, these findings illustrate how the host’s metabolic state influences *S. indica* colonization and highlight the importance of future research on the impact of nutrient availability during symbiotic interactions. Additionally, our results indicate that the growth phenotype can occur independently of hypercolonization and cell death, emphasizing distinct regulatory pathways for these processes.

Our data show that autophagy activation is involved in restriction of fungal colonization and is important to promote cell survival during dAdo-induced cell death. We propose that the activation of autophagy in colonized cells, promotes a protective response to limit fungal-induced stress and cell death (Fig. 5). These results also emphasize the complexity of the *S. indica* colonization process, in which nutritional factors, metabolic status and immune-related cell death are modulated by autophagy.

## Materials and Methods

### Plant lines

*Arabidopsis thaliana* ecotype Columbia (Col-0) was used as a wild-type (WT) control. The T-DNA insertion mutant lines in this study are: *atg5-1* (AT5G17290) SAIL-129B07, *atg5-3* SALK-020601C (Thompson et al., 2005; Have et al., 2019), *atg10-1* (AT3G07525) SALK-084434, *atg11-1* (AT4G30790) SAIL-1166G10, *atg2-2* (AT3G19190) EMS-mutant (Wang et al., 2011), *ent3* (AT4G05120) SALK-204257C. The transgenic lines *pUbi::mCherry-ATG8E* in Col-0 WT and *atg5-1* background (Stephani et al., 2020). The double mutant *atg5 ent3* was obtained by crossing *atg5-3* and *ent3* lines. All transgenic lines were genotyped by PCR and homozygous lines were isolated.

### Plant-growth conditions

Surface-sterilized seeds of Arabidopsis were germinated and grown on ½ MS (Murashige-Skoog Medium, with vitamins, pH 5.7) containing 0.5% (w/v) sucrose and 0.4% (w/v) Gelrite (Duchefa, Haarlem, the Netherlands) and stratified in darkness for 3 days at 4◦C. Plants were grown under short day conditions (8 h light, 16 h dark) with 130 µmol m^−2^ s^−1^ of light and 22°C /18°C.

### Fungal strains and culturing techniques

*Serendipita indica* strain DSM11827 (German Collection of Microorganisms and Cell Cultures) Braunschweig, Germany) was grown on complete medium (CM) containing 2% (w/v) glucose and 1.5% (w/v) agar at 28*^◦^*C as described (Hilbert et al., 2012). For confocal microscopy studies, the *S. indica* strain constitutively expressing a *S. indica* codon-optimized GFP gene was used (Hilbert et al., 2012).

### Fungal inoculation

Eight-day-old Arabidopsis seedlings were transferred to ½ MS (Murashige-Skoog Medium, with vitamins, pH 5.7) and 0.4% (w/v) Gelrite plates. Nine-day-old seedlings, specifically the roots and the surrounding area, were inoculated with 800 µL containing 5×10^5^ *S. indica* chlamydospores / mL. Control plants were inoculated with sterile milli-Q water as mock treatment. At the indicated time points, a 4 cm root section was harvested starting 0.5 cm below the shoot. Colonized roots were washed thoroughly to remove extraradical hyphae and frozen in liquid nitrogen. Each biological replicate contains three plates with 15 seedlings each. For seed inoculation, surface-sterilized Arabidopsis seeds were incubated in 1 mL containing 5×10^5^ *S. indica* chlamydospores / mL or sterile milli-Q water for 1 hour followed by their distribution on ½ MS and 0.4% (w/v) Gelrite plates.

### RNA extraction and real-time quantitative PCR analysis

Total RNA was extracted from colonized or mock-treated ground plant root material using TRIzol reagent (Invitrogen, Thermo Fisher Scientific, Schwerte, Germany) according to the manufacturer’s instructions and digested with DNase I to prevent genomic DNA contamination (Thermo Fisher Scientific, Schwerte, Germany). cDNA was synthesized with 1 µg total RNA primed with Oligo-dT and random hexamers primers using the First Strand cDNA Synthesis Kit (Thermo Fisher Scientific, Schwerte, Germany). Quantitative real-time PCR was performed using the 2x GoTaq qPCR master mix (Promega, Walldorf, Germany) with 10 ng of cDNA template and 0.5 µM of each oligonucleotide in a final volume of 15 µL. Reactions were amplified in a CFX connect real time system (BioRad, Munich, Germany) according to the following protocol 95*^◦^*C 3 min, 95*^◦^*C 15 s, 59*^◦^*C 20 s, 72*^◦^*C 30 s, 40 cycles and a melting curve analysis. Relative expression was calculated using the 2^-ΔCt^ method. Sequences of all primers can be found in Table S1.

### PAM fluorometric measurements

For chlorophyll fluorescence analysis, eight-day-old Arabidopsis seedlings were transferred to 24-well plates (three per well) containing 2 mL of 2.5 mM MES buffer (pH 5.6). After overnight regeneration, seedlings were treated with mock (MES 2.5 mM buffer), dAdo or methyl jasmonate (Sigma-Aldrich, Taufkirchen, Germany) to a final concentration of 500 µM. Maximum quantum yield of photosystem (PS)-II (F_V_/F_M_) of dark-adapted samples was quantified using Pulse Amplitude Modulation (PAM) fluorometry (M-Series PAM fluorometer, Heinz Walz GmbH, Effeltrich, Germany). Data were analyzed using ImagingWin software (V2.56p; Walz, Germany).

### Seed germination assays

Surface-sterilized seeds of Arabidopsis were transferred to 24 well plates, containing 2 mL of 1/10 PNM (Plant Nutrition Medium, pH 5.6). The medium was supplemented with 500 µM dAdo or mock (2.5 mM MES, pH 5.6). 10 seeds were placed into each well and after 2 days of stratification, grown under short day conditions. The growth of the seedlings was monitored via PAM fluorometry at 7, 14 and 21 days after transfer to the growth chamber. Analysis of the photosynthetic active area was analyzed using the software Fiji (ImageJ).

### Root length measurements

Surface-sterilized Arabidopsis seeds were inoculated with mock or *S. indica* spores on plates containing 1/10 PNM (Plant Nutrition Medium, pH 5.7). 21 days after inoculation, scans of the square plates containing seedlings were taken. Images were analyzed using Fiji (ImageJ) to measure the length of the primary root (cm) of seedlings developing true leaves.

### Extraradical colonization assays

Quantification of extraradical colonization of *S. indica* on Arabidopsis was performed on seed-inoculated plants grown for 12 days and stained with the chitin-binding lectin stain Wheat Germ Agglutinin conjugated with Alexa Fluor 488 (WGA-AF 488, Invitrogen Thermo Fisher Scientific, Schwerte, Germany). Mock and *S. indica*-treated seedlings were stained directly on plate with 1x PBS solution containing WGA-AF 488 (5 µL / mL from 1 mg/mL stock solution) and incubated for 1-2 minutes. Subsequently, the roots were washed with 1x PBS solution. The stained seedlings were transferred to a new fresh plate and fluorescence detection was conducted using an Odyssey M Imaging System (LI-COR Biosciences). WGA-AF 488 fluorescence intensity was quantified using ImageJ by subtracting background signal and normalizing to the root length of the different genotypes.

### Cell death staining with Evans Blue

Root cell death was quantified using Evans blue dye (Sigma-Aldrich, Taufkirchen, Germany) to assess *S. indica*- or dAdo-induced cell death at the indicated time points. Treated roots were washed three times with milli-Q water before staining with 2.5 mM Evans blue solution in 0.1 M CaCl_2_ pH 5.6 for 15 min, based on a modified version of the protocol described by Vijayaraghavareddy et al., 2017. After 1 hour of washing, root images were taken using a Leica M165 FC stereo microscope. The microscopy of *S. indica* colonized samples was performed using the differentiation zone of the Arabidopsis roots, and the microscopy of the roots treated with dAdo corresponds to the root tip. Quantification of cell death was performed using Fiji (ImageJ).

### Autophagic flux assay

For carbon starvation treatment, Arabidopsis seedlings expressing *pUbi::mCherry-ATG8E* in WT or in the *atg5-1* KO mutant background were grown on ½ MS containing 1% (w/v) sucrose and 0.4% (w/v) Gelrite. After 7 days, seedlings were transferred to ½ MS without sucrose and plates were covered with aluminium foil and grown under the same conditions for 9 days (Stephani et al., 2020). Whole seedlings were harvested and frozen in liquid nitrogen. For analysis during *S. indica* colonization, surface-sterilized Arabidopsis seeds were inoculated with mock or *S. indica* treatment. After 10 days, root tissue was harvested and frozen in liquid nitrogen.

### Protein extraction and immunoblot analysis

Frozen samples were homogenized with a bead mill (TissueLyser II, Qiagen) for 2 min (frequency 30^-1^). 1x lysis buffer (2x lysis buffer: 300 mM NaCl, 100 mM HEPES pH 7.4, 2 mM EDTA, 2% Triton X-100) with 1x plant protease inhibitor cocktail (Sigma-Aldrich, Taufkirchen, Germany) and 1 mM PMSF was added, and samples were vortexed. The lysates were cleared by centrifugation at 13,000 g for 10 min at 4 °C. Protein concentration was determined using the Pierce BCA Protein Assay Kit (Thermo Fisher Scientific, Schwerte, Germany). The protein extract was mixed with Laemmli buffer (6x) containing beta-mercaptoethanol and then boiled for 10 min at 95°C. SDS-PAGE was performed with 10% gels and 1x Tris Glycine-SDS buffer (BioRad, Munich, Germany). Semi-dry blotting was performed on PVDF membranes. Membranes were blocked with 3% BSA (VWR, Darmstadt, Germany) in TBS and 0.05% Tween 20% (v/v) (TBS-T) for 1 hour at room temperature. After incubation overnight at 4◦C with the primary antibody anti-mCherry (1:1000, 5993, BioVision) diluted in 1x phosphate-buffered saline (PBS), the membranes were washed three times with TBS-T. After a 40 min incubation with the secondary antibodies IRDye 680RD / 800CW (1:10,000, LI-COR) diluted in 3% BSA TBS-T, the membranes were washed three times with TBS-T and finally rinsed with 1x PBS. Fluorescent Western blot detection was performed using an Odissey DLx (LI-COR Biosciences GmbH, Bad Homburg vor der Höhe, Germany).

### Confocal imaging and image quantification

A TCS SP8 confocal microscope (Leica, Wetzlar, Germany) was used for confocal laser scanning microscopy on living cells. mCherry was excited by a laser light at 561 nm and the emitted light was detected with a hybrid detector (HyD2) at 602-638 nm. The mCherry-ATG8-labelled punctate structures were counted in a 63x captured image size (184.52 µm × 184.52 µm) considering maximal projections of 8-10 frames with 4 µm step size.

### Statistical analysis

Analysis was performed using GraphPad Prism software (v.9.4.1 for Windows) or RStudio (R v.4.1.1 for Windows). The detailed statistical method is given in the figure legends.

### Transcriptomic analysis (RNA sequencing)

Arabidopsis Col-0 WT roots were inoculated with mock (milli-Q water) or *S. indica*. Roots were harvested at 1, 3, 6, and 10 days post-inoculation. For each condition three biological replicates were considered. For each sample, stranded RNA-Seq libraries were generated and quantified by qPCR (Eichfeld et al., 2023). The RNA-Seq libraries were generated and sequenced at US Department of Energy Joint Genome Institute (JGI) under a project proposal (Proposal ID: 505829) (Zuccaro & Langen, 2020). The raw reads were filtered and trimmed using the JGI QC pipeline. Filtered reads from each library were aligned to the Arabidopsis (TAIR10) reference genome using HISAT2 (Kim et al., 2015) and the gene counts were generated using featureCounts (Liao et al., 2014). DESeq2 was used to perform differential gene expression analysis (Love et al., 2014). Autophagy-associated genes were identified using the Gene ontology (GO) terms annotated by Denny et al. (2018) and retrieved from the Arabidopsis (TAIR10) annotation version 2023-05-01.

## Acknowledgments

We would like to thank Prof. Ralph Hückelhoven and Prof. Martin Stegmann for providing constructive feedback. We would like to thank Lisa Mahdi for conducting experiments for the RNA-seq analysis. We extend our gratitude to the U.S. Department of Energy, Joint Genome Institute (https://ror.org/04xm1d337) and Yu Zhang, Sravanthi Tejomurthula, Daniel Peterson, Vivian Ng & Igor Grigoriev for producing sequencing data within the work proposal 10.46936/10.25585/60001292.

## Author contributions

P.Z.R. and A.Z. conceived and designed the study, analyzed the data, and wrote the manuscript. P.Z.R., E.L., N.C., N.D., A.M., performed the experiments and analyzed the data. C.D.Q. performed bioinformatics analysis and figures. E.L., N.C., N.D., G.L., C.D.Q., and Y.D., contributed to the design of this research and reviewed and edited the manuscript. A.Z. supervised the research and provided resources, laboratory infrastructure and funding. All authors read and approved the final manuscript.

