## Supplemental Figures for "Autophagy restricts fungal accommodation in the roots of *Arabidopsis thaliana*"

### Supplementary figures

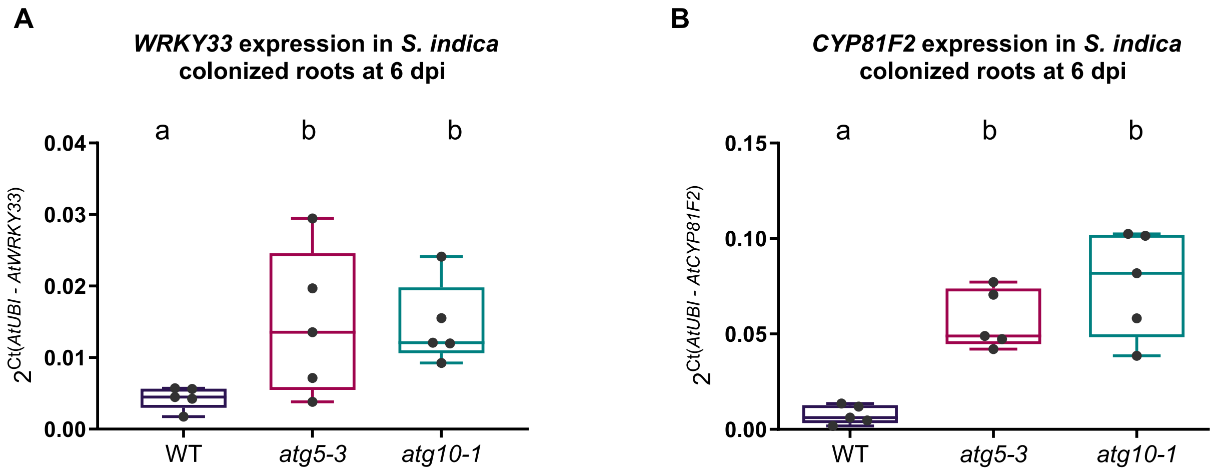


**Figure S1: Expression of immune marker genes in *A. thaliana* WT and autophagy mutants *atg5-3* and *atg10-1*.**

(**A-B**) Expression of *WRKY33* and *CYP81F2* marker genes in WT and autophagy mutants *atg5-3* and *atg10-1* from *S. indica*-colonized roots at 6 dpi. Relative expression was calculated compared to plant (*AtUbi*) using cDNA as template and the $2^{-\Delta Ct}$ method. Boxplots with whiskers extending to the minimum and maximum values represent data from 5 independent biological replicates. Different letters indicate significant differences ($p < 0.05$) according to Kruskal-Wallis test and post-hoc Dunn test using Benjamini-Hochberg for false discovery rate correction.

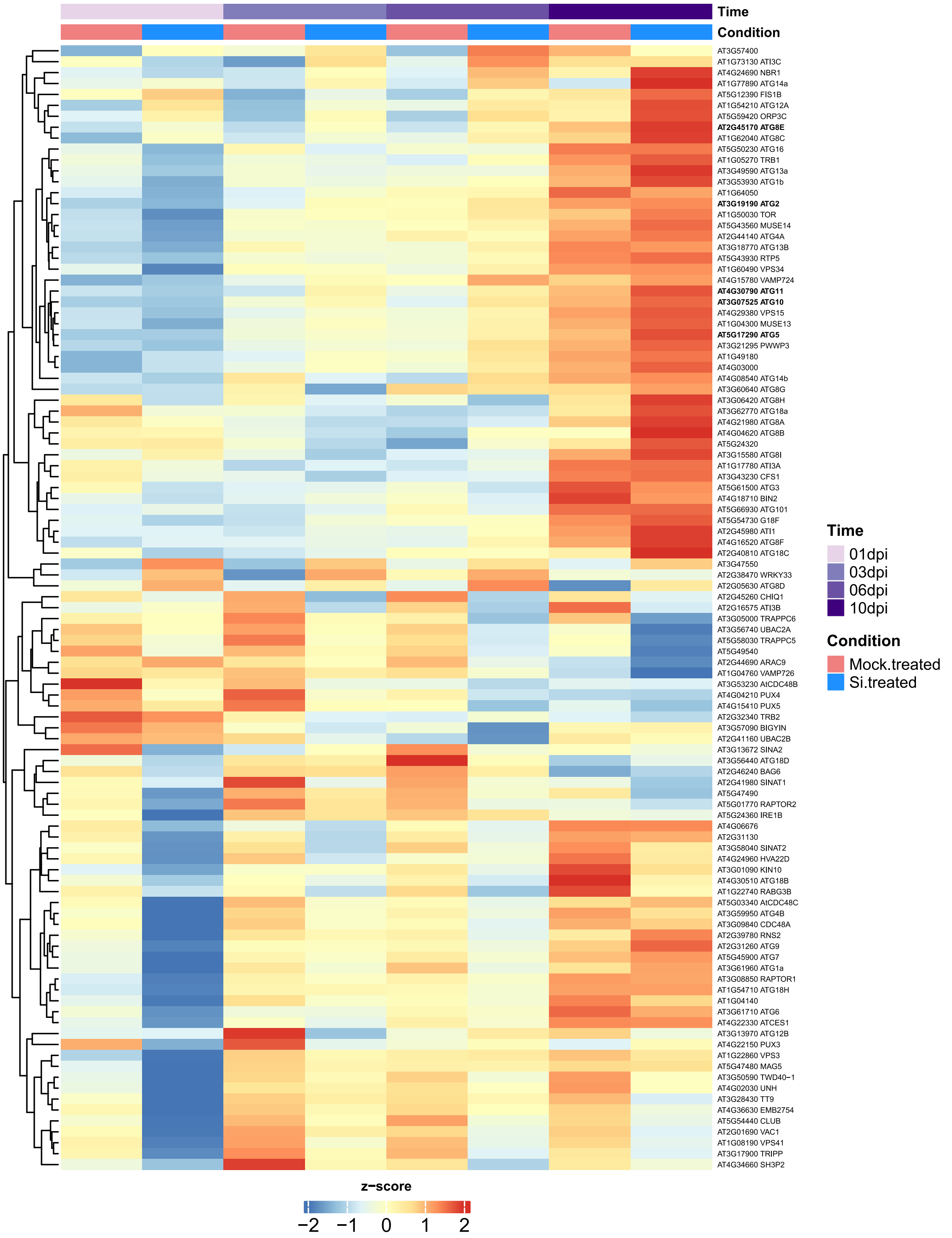


**Figure S2: Expression of autophagy-associated genes during *S. indica* colonization.**

The heatmap shows expression level of autophagy-associated genes in Arabidopsis WT roots samples collected at 1, 3, 6 and 10 days post inoculation with S. *indica* (*Si*) or mock. Genes with an average TPM value > 1 TPM across all samples were selected. The heatmap shows the z-score of log2 transformed TPM + 1 values of selected Arabidopsis genes. For each condition the average expression across the three biological replicates is represented. The results of DESeq2 analysis from pairwise comparisons (*S. indica*-treated vs mock-treated samples) at each timepoint are in Table S2. (Zuccaro, A.,& Langen, G. 2020; Eichfeld et al., 2023).

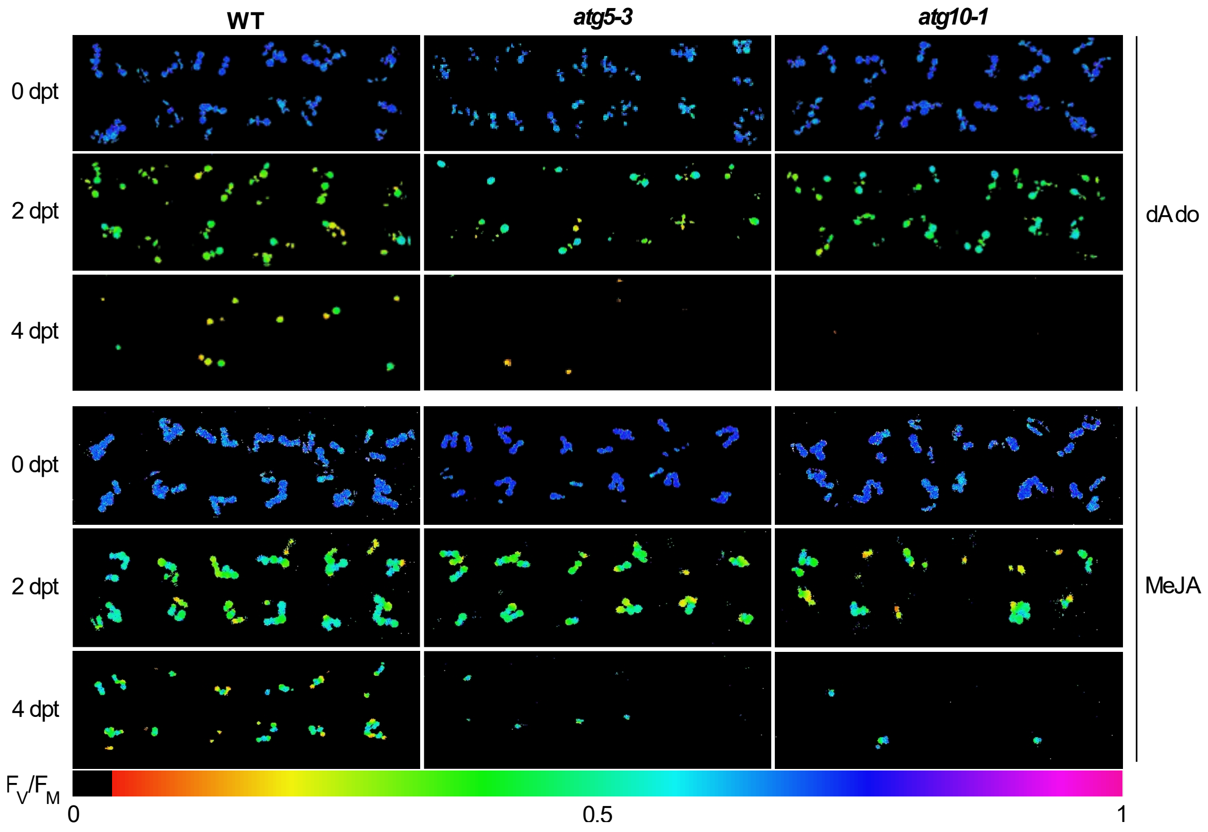


**Figure S3: Autophagy mutants *atg5-3* and *atg10-1* show increased sensitivity upon dAdo treatment.**

Visualization of photosystem II maximum quantum yield ($F_V/F_M$) of WT and autophagy mutants *atg5-3* and *atg10-1* after dAdo or MeJA (500 µM) at 0, 2 and 4 days post treatment. The $F_V/F_M$ value is illustrated by the color scale shown below. In each treatment each of the 12 wells contain three seedlings.

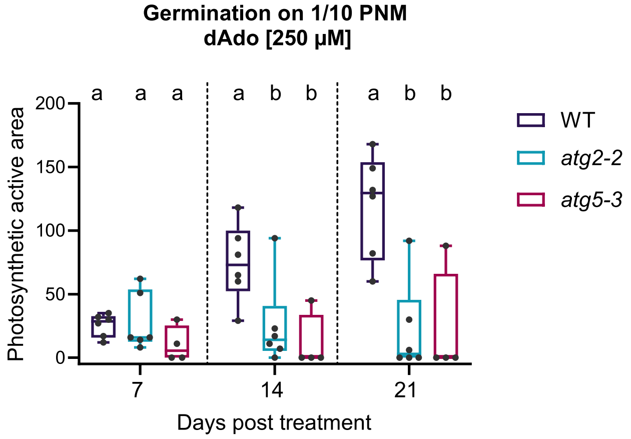


**Figure S4: Seed germination assay in *A. thaliana* WT and autophagy mutants *atg5-3* and *atg2-2*.**

Quantification of photosynthetic active area after 7, 14 and 21 days post treatment. Surface-sterilized seeds were treated with mock (MES 2.5 mM buffer) or dAdo (500 µM). Boxplots with whiskers extending to the minimum and maximum values represent relative values from 6 independent biological replicates. Different letters indicate significant differences ($p < 0.05$) according to one-way ANOVA with post-hoc Tukey HSD test.

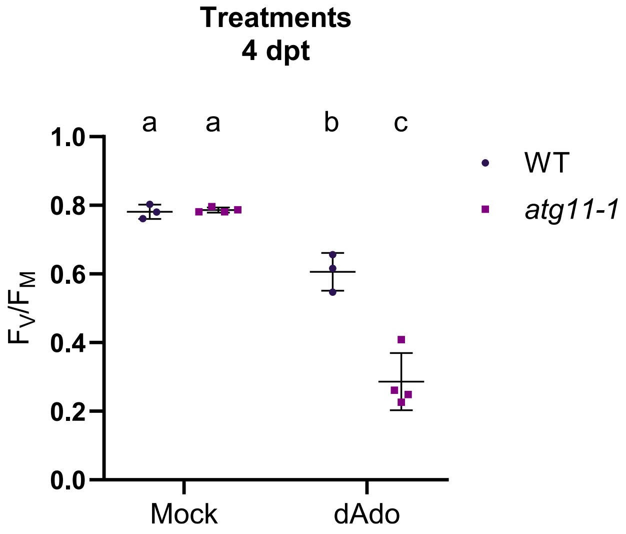


**Figure S5: Autophagy mutant *atg11-1* show increased sensitivity upon dAdo treatment.**

Quantification of photosystem II maximum quantum yield ($F_V/F_M$) of WT and *atg11-1* at 4 dpt by PAM fluorometry. Nine-day-old seedlings were treated with mock (MES 2.5 mM buffer) or dAdo (500 µM). The plot (mean ± SD) represents data from 3-4 independent biological replicates, each consisting of 12 wells with 3 seedlings per well. Different letters indicate significant differences ($p < 0.05$) according to one-way ANOVA with post-hoc Tukey HSD test.

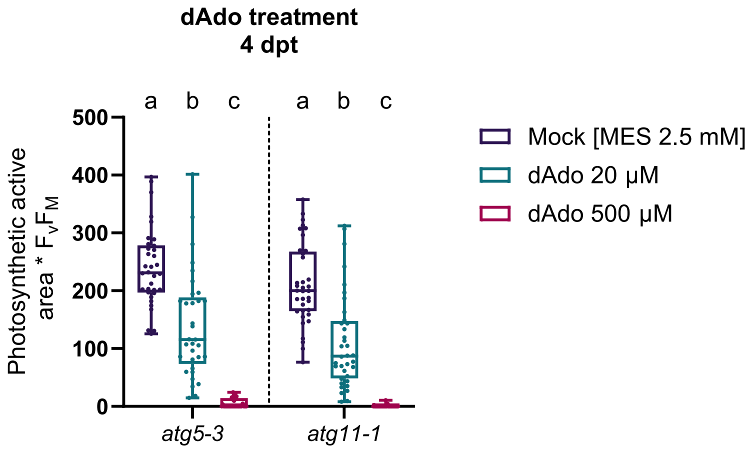


**Figure S6: dAdo dose-dependent response on autophagy mutants.**

Quantification of photosystem II maximum quantum yield ($F_V/F_M$) of *atg5-3* and *atg11-1* at 4 dpt by PAM fluorometry. Nine-day-old seedlings were treated with mock (MES 2.5 mM buffer), dAdo (250 µM) or dAdo (500 µM). Boxplots with whiskers extending to the minimum and maximum values represent relative values from 33-36 biological replicates. Different letters indicate significant differences ($p < 0.05$) according to one-way ANOVA with post-hoc Tukey HSD test.

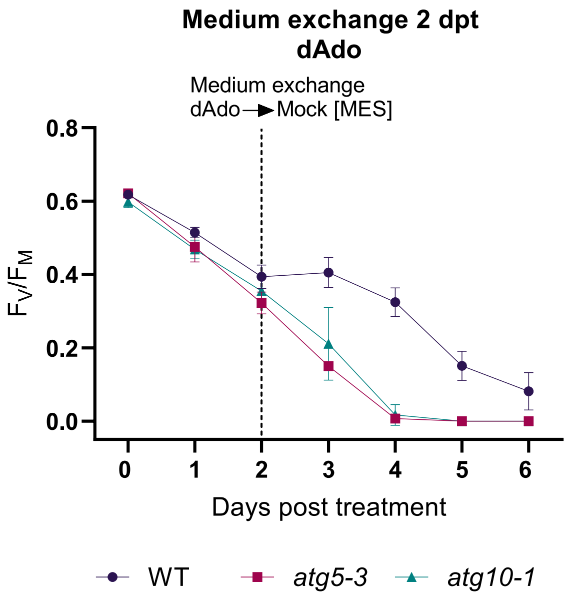


**Figure S7: dAdo recovery assay in *A. thaliana* WT and autophagy mutants *atg5-3* and *atg10-1*.**

Photosystem II maximum quantum yield ($F_V/F_M$) of nine-day-old seedlings treated with mock (MES 2.5 mM buffer) or dAdo (500 µM), measured by PAM fluorometry. dAdo treatment solution was replaced after 48 h with MES 2.5 mM buffer. Error bars represent ± SD of the mean of 3 independent biological replicates, each consisting of 12 wells with 3 seedlings per well.

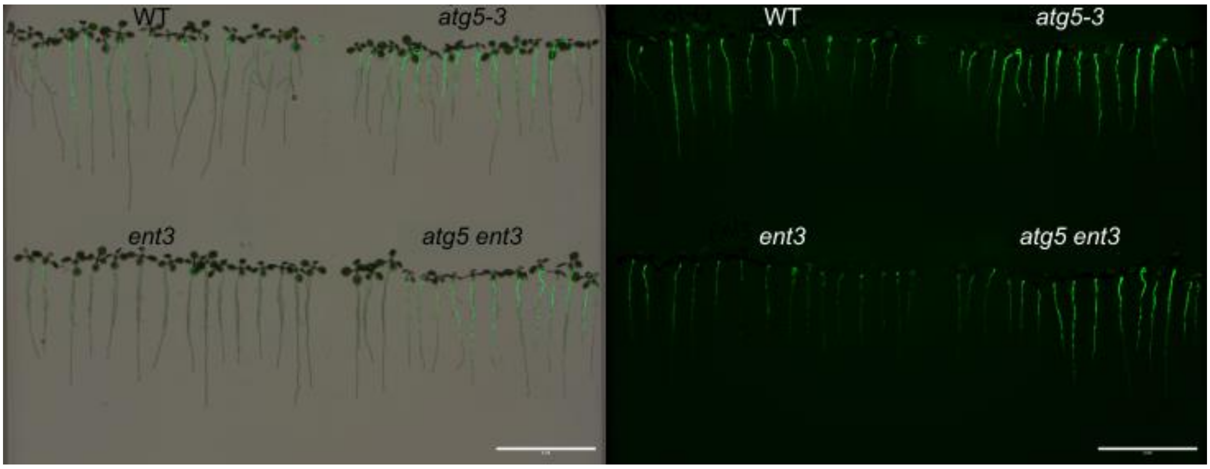


**Figure S8: Extraradical *S. indica* colonization of 12-days Arabidopsis seedlings**

Representative image of seed-inoculated Arabidopsis seedlings stained with WGA-AF 488 for *S. indica* detection. Fluorescence was detected using an Odyssey M Imaging System. Left and right images display the roots captured in the bright field channel and Alexa Fluor 488 channel, respectively. Scale bar: 4 cm.

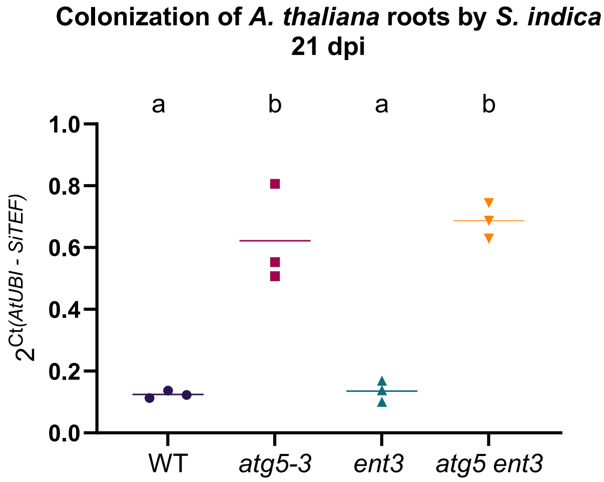


**Figure S9: Colonization by *S. indica* on 1/10 PNM after 21 days**

Quantification of endophytic colonization by RT-qPCR of *S. indica* seed-inoculated plants grown for 21 days. Fungal colonization in plant root tissue was calculated from the ratio of S. indica (*SiTEF*) to plant (*AtUbi*) using cDNA as template and the $2^{-\Delta Ct}$ method. The plot (mean ± SD) represents data from 3 independent biological replicates. Different letters indicate significant differences ($p < 0.05$) according to one-way ANOVA with post-hoc Tukey HSD test.
